# Acetylcholine release inhibits distinct excitatory inputs onto hippocampal CA1 pyramidal neurons via different cellular and network mechanisms

**DOI:** 10.1101/575811

**Authors:** Priyodarshan Goswamee, A. Rory McQuiston

## Abstract

In hippocampal CA1, muscarinic acetylcholine (ACh) receptor (mAChR) activation via exogenous application of cholinergic agonists has been shown to presynaptically inhibit Schaffer collateral (SC) glutamatergic inputs in stratum radiatum (SR), and temporoammonic (TA) and thalamic nucleus reuniens (RE) glutamatergic inputs in stratum lacunosum-moleculare (SLM). However, steady-state uniform mAChR activation may not mimic the effect of ACh release in an intact hippocampal network. To more accurately examine the effect of ACh release on glutamatergic synaptic efficacy, we measured electrically evoked synaptic responses in CA1 pyramidal cells (PCs) following the optogenetic release of ACh in genetically modified mouse brain slices. The ratio of synaptic amplitudes in response to paired-pulse SR stimulation (stimulus 2/stimulus 1) was significantly reduced by the optogenetic release of ACh, consistent with a postsynaptic decrease in synaptic efficacy. The effect of ACh release was blocked by the M_3_ receptor antagonist 4-DAMP, the GABA_B_ receptor antagonist CGP 52432, inclusion of GDP-β-S, cesium, QX314 in the intracellular patch clamp solution, or extracellular barium. These observations suggest that ACh release decreased SC synaptic transmission through an M_3_ muscarinic receptor-mediated increase in inhibitory interneuron excitability, which activate GABA_B_ receptors and inwardly rectifying potassium channels on CA1 pyramidal cells. In contrast, the ratio of synaptic amplitudes in response to paired-pulse stimulation in the SLM was increased by ACh release, consistent with presynaptic inhibition. ACh-mediated effects in SLM were blocked by the M_2_ receptor antagonist AF-DX 116, presumably located on presynaptic terminals. Therefore, our data indicate that ACh release differentially modulates excitatory inputs in SR and SLM of CA1 through different cellular and network mechanisms.

## Introduction

Medial septum and diagonal band of Broca complex (MS/DBB) cholinergic neurons project to the hippocampus where they influence attention, learning and memory (Hasselmo 2006), and synaptic plasticity (Zhenglin Gu and Yakel 2011). In Alzheimer’s disease, MS/DBB cholinergic neurons are among the first cells to degenerate (Hampel et al. 2018), which underscores their significance in the pathophysiology of the disease and highlights a need for a better understanding of the mechanisms by which MS/DBB cholinergic inputs affect hippocampal function.

MS/DBB cholinergic inputs affect hippocampal function through a variety of cellular and network mechanisms. In hippocampal CA1, activation of muscarinic and nicotinic receptors affect the excitability of pyramidal cells (PCs) (Cole and Nicoll 1983), inhibitory interneurons (Frazier et al., n.d.; Jones and Yakel 1997; McQuiston and Madison 1999b, 1999c, 1999a; Pitler and Alger 1992) and astrocytes (Araque et al. 2002). At the synapse, presynaptic nicotinic receptor activation has been shown to facilitate the release of glutamate onto CA1 PCs (Ji et al, 2001; Maggi et al, 2003; Sola et al, 2006) whereas activation of presynaptic muscarinic receptor has been shown to inhibit glutamate release (M. Hasselmo and Schnell 1994; Qian and Saggau 1997; Valentino and Dingledine 1981).

Hippocampal CA1 PCs are primarily driven by three excitatory glutamatergic inputs. Two of these excitatory inputs are located in the stratum lacunosum-moleculare and synapse on distal apical CA1 PC dendrites - the temporoammonic pathway (TA) from the entorhinal cortex and inputs from the thalamic nucleus reuniens (RE). The other excitatory input, the Schaffer-collaterals (SC), originate in hippocampal CA3 and synapse on the proximal apical and basal dendrites of CA1 PCs in the stratum radiatum (SR) and stratum oriens (SO), respectively. A proportionately larger presynaptic muscarinic receptor-mediated inhibition has been observed in the SC pathway of SR compared to inhibition of inputs located in the SLM (M. Hasselmo and Schnell 1994). However, these studies were conducted using exogenous uniform activation of muscarinic receptors in rat brain slices. This experimental paradigm may not accurately measure the influence of cholinergic inputs on hippocampal CA1 synaptic function as MS/DBB cholinergic inputs and acetylcholinesterase are not uniformly distributed in the hippocampus (Aznavour et al., 2002; Franklin and Paxinos, 2007). Physiological measures of acetylcholine levels in the hippocampus match the anatomical data as acetylcholine concentrations appear to be larger in the stratum pyramidale compared to all other layers in hippocampal CA1 during theta rhythm (Zhang et al., 2010). Thus, muscarinic receptors located on different subsets of presynaptic glutamatergic terminals may be exposed to different concentrations of acetylcholine following release from MS/DBB terminals. This would impact the magnitude of cholinergic presynaptic inhibition following the release of ACh from synaptic terminals.

Therefore, we assessed the impact of ACh release on glutamatergic synaptic transmission in SR and SLM of hippocampal CA1. To do this we optogenetically released ACh from MS/DBB cholinergic terminals expressing the red-shifted optogenetic excitatory protein ReaChR and measured the effect of ACh release on paired electrically-evoked glutamatergic postsynaptic excitatory responses in hippocampal CA1 PCs. Here, we report that optogenetic activation of ACh release indirectly reduced paired-pulse ratio (PPR, stimulus 2/stimulus 1) in SR by an M_3_-muscarinic receptor-mediated increase in excitability of inhibitory interneurons. The combined increase in interneuron excitability coupled with SR excitatory inputs resulted in the activation of postsynaptic GABA_B_-receptors and inwardly rectifying potassium channels in the CA1 PCs. In contrast, ACh release appeared to result in presynaptic inhibition of terminals in SLM via an M_2_-dependent mechanism. Therefore, our data suggest that ACh release has different cellular and network mechanisms of action on glutamatergic neurotransmission in the SR and SLM of hippocampal CA1.

## Materials & Methods

### Pharmacological agents

All chemicals were purchased from ThermoFisher scientific unless otherwise indicated. VU 0255035 (highly selective muscarinic M_1_ receptor antagonist), 4-DAMP (Muscarinic M3 receptor antagonist), PD 102807 (selective M4 receptor antagonist), AF-DX 116 (selective M2-muscarinic receptor antagonist), Baclofen (GABA_B_ receptor antagonist), QX 314 chloride (intracellular sodium channel blocker), and CGP 52432 (selective GABAB receptor antagonist) were obtained from Tocris Bioscience (Ellisville, Missouri). GDP-β-S was purchased from Sigma. Bicuculline methochloride (competitive GABA_A_ receptor antagonist) was purchased from helloBio (Montogomery, New Jersey). Biocytin (B-1592) was purchased from Life Technologies (Invitrogen).

### Animals

The B6.Cg-Gt(ROSA)26Sortm2.2Ksvo/J (ReaChR JAX Stock No. 026294) and 134 B6; 129S6-Chattm1(cre)Lowl/J (Chat-cre, JAX Stock No. 006410) mice used in these studies were housed in an animal care facility approved by the American Association for the Accreditation of Laboratory Animal Care (AAALAC). Animal experimental procedures followed a protocol approved by the Institutional Animal Care and Use Committee of Virginia Commonwealth University (AD 20205). This protocol adhered to the ethical guidelines described in The Care and Use of Laboratory Animals 8th Edition. Efforts were made to minimize animal suffering and to reduce the number of animals used.

### Breeding Strategy

Homozygous chat-cre mice were crossed with homozygous ReaChR mice. The resulting offspring were heterozygous for both alleles. In some experiments, these progeny mice, which were heterozygous for both alleles, were further crossed to achieve homozygosity in both alleles. Animals that expressed at least one copy of the mutant alleles in both loci were utilized for physiological experiments and were identified by genotyping using specific primers for the mutant alleles (Table1).

**Table 1:**
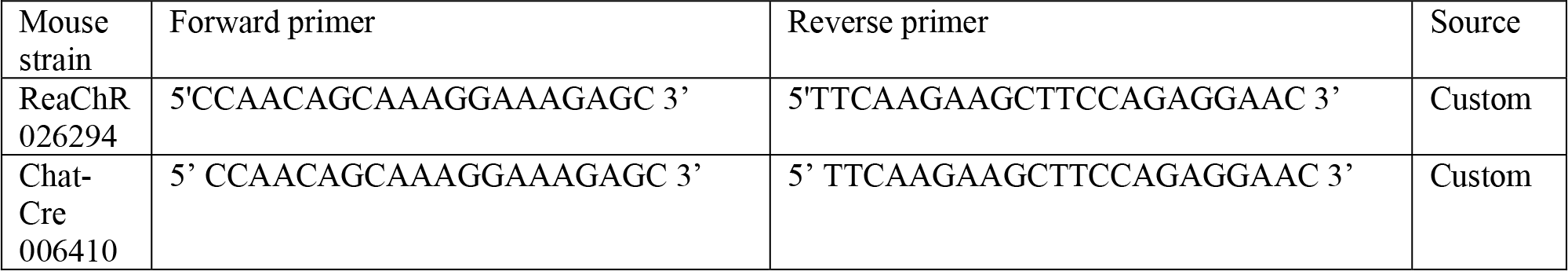
PCR primers used to genotype mouse crosses. In brain slices prepared from some animals, ectopic expression of mCitrine was observed in what appeared to be astrocytes. Data from such animals were not included in analysis.

### Preparation of brain slices

Adult mice (30 to 180 days old) were deeply anesthetized with an intraperitoneal injection of ketamine (200 mg/kg) and xylazine (20 mg/kg), following which, the animals were transcardially perfused with ice-cold sucrose saline (consisting of (in mM): Sucrose 230, KCl 3.0, CaCl_2_ 1.2, MgCl_2_ 6, Na_2_HPO_4_ 1.2, NaHCO_3_ 25, glucose 25). The brain was removed and sectioned to prepare coronal MS/DBB or horizontal brain slices that contained mid-temporal hippocampus. Slices were cut at 350 μm on a Leica VT1200 (Leica Microsystems, Buffalo Grove, IL, USA) and incubated in a holding chamber containing artificial cerebrospinal fluid (aCSF) ((in mM): NaCl 125, KCl 3.0, CaCl_2_ 1.2, MgCl_2_ 1.2, NaHPO_4_ 1.2, NaHCO_3_ 25, glucose 25 bubbled with 95% O_2_ / 5% CO_2_) at 32°C for at least 30 min before commencement of electrophysiological experiments.

### Light-evoked release of acetylcholine from MS/DBB cholinergic axon terminals

Cholinergic terminals expressing ReaChR-mCitrine were stimulated by 40 yellow light pulses delivered at 4-5 Hz (10 ms in duration). The light pulses were generated from a UHP-T-LED-White light-emitting diode (LED) (Prizmatix Modiin-Ilite, Givat Shmuel, Israel). The white light exiting the LED was filtered by an HQ 550-600/50x excitation filter and was focused into the epi-illumination light path of the Olympus BX51WI microscope and back aperture of a 10x water immersion objective (0.3 NA) using an optiblock beam combiner (Prizmatix) and a dichroic mirror (700dcxxr, Chroma Technology) in the filter turret.

### Immunohistological Verification of ReaChR Expression in Septal Cholinergic Neurons

Coronal MS/DBB sections containing one biocytin-filled mCitrine expressing cell were drop-fixed in 4% paraformaldehyde for at least 24 hours. Subsequently, slices were washed and incubated in a blocking/permeabilizing buffer (1X PBS supplemented with 0.2% bovine serum albumin and Triton-X 100) for 24 h. Sections were then incubated for 3 days at 4 °C with antibody cocktail comprising of 1:100 dilution of a Goat polyclonal anti-Chat antibody (EMD Millipore, Cat# AB144P) and 1:200 dilution of GFP-Tag polyclonal antibody conjugated with AlexaFluor 488. Slices were then washed 3 times with phosphate-buffered saline and incubated with 1:200 dilution of Donkey anti-Goat 568 (Thermo Fisher, Cat # A-11057) and 1:1000 dilution of streptavidin Alexa Fluor 633 (Thermo Fisher, Cat # S-11226). Processed slices were then imaged using a Zeiss LSM710 confocal microscope (Carl Zeiss, Jena, Germany).

### Electrophysiology

Whole cell patch clamp recordings were conducted on medial septum/diagonal band of Broca (MS/DBB) cholinergic neurons, hippocampal CA1 interneurons, and PCs. For these experiments, patch pipettes (3-4 MΩ) pulled from borosilicate glass (8250 1.65/1.0 mm) on a Sutter P-1000 pipette puller and were filled with intracellular recording solution that contained either a potassium-based recording solution ((in mM): KMeSO_4_ 145, NaCl 8, Mg-ATP 2, Na-GTP 0.1, HEPES 10, EGTA 0.1) or a Cesium-based recording solution ((in mM): CsMeSO_4_ 120, NaCl 8, Mg-ATP 2, Na-GTP 0.1, HEPES 10, Cs-BAPTA 10, QX-314 Chloride 10). In some experiments with the potassium recording solution, the GTP was replaced with 5μM GDP-β-S, an inhibitor of G-protein coupled receptor. 0.1% biocytin was included in the intracellular recording solution in a subset of experiments for post hoc identification of the recorded cell. Membrane potentials or excitatory postsynaptic currents (EPSCs) were measured with a Model 2400 patch clamp amplifier (A-M Systems, Port Angeles, WA) and converted into a digital signal by a PCI-6040E A/D board (National Instruments, Austin, TX). WCP Strathclyde Software (courtesy of Dr. J Dempster, Strathclyde University, Glasgow, Scotland) was used to collect and store membrane potential or EPSC responses on a PC computer. For all voltage clamp experiments, series resistance was compensated to approximately 70%, and experiments in which the access changed by more than approximately 20% was discarded. To evoke paired-pulse responses in hippocampal CA1 principal cells, bipolar platinum-iridium stimulating electrodes (approx. 100 kΩ, FHC Inc., Bowdoin, ME, USA) were placed in SC (in stratum radiatum) or TA/RE pathway (stratum lacunosum-moleculare). A pair of electrical pulses (40-120 μs pulse width, 50 −100 μAmp) 50 ms interval was utilized to stimulate the axons. Stimulation currents were titrated to achieve responses that were approximately 70% of the maximum EPSC amplitudes. The time interval between successive paired-pulse stimulation was 45 s. The PPR was calculated by dividing the averaged peak EPSC amplitude in response to the second pulse (P2) by that of the first pulse (P1) (P2/P1). Light-evoked ACh release was achieved by delivering a 5 Hz train of 40 yellow light pulses (10 ms each) immediately before the electrical paired-pulse. This stimulation paradigm will be referred to as “Light ON” for the remainder of the manuscript. In the same cells, PPR was calculated in the absence of light stimulation. These responses will be termed as “Light OFF”. Data presented shows averaged responses from 5 consecutive Light OFF and Light ON stimulations. Because optogenetic release of ACh in slices is prone to rapid hydrolyzation due to acetylcholinesterase activity, we conducted some experiments in presence of 0.1 uM Donepezil, an acetylcholinesterase inhibitor, as utilized by others (Alger, Nagode, and Tang 2014).

### Experimental Design and Statistical Analysis

Data analysis was performed with OriginPro 2018 (OriginLab Corp., Northampton, MA, USA) and Excel (Microsoft, Redmond, WA). Statistics were performed using GraphPad Prizm software (La Jolla, California USA). A total of 39 mice (21 males and 18 females) were utilized in the study. At least 3 animals were utilized for every experiment. Numbers of paired recording performed for every experiment was aimed to attain 80% power, determined using GraphPad StatMate 2.0 (San Diego, California, USA). The clustering of cells per mice was random, and no data point was utilized in more than one analysis. The precise number of cells per animal sampled is indicated in the results. Comparison of PPR amplitudes were conducted within cell and had only one variable (*i.e.*, light-evoked release of ACh) and therefore, the statistical significances of the results were determined using paired 2-tailed t-tests. Statistical differences of results between two groups of data collected from separate groups of cells (Figs. 4E and 6A) were compared using 2 tailed unpaired t tests. Differences were determined to be statistically significant for p-values less than 0.05. All data are reported as the mean, standard error of the mean (SEM). Asterisks were used as follows unless otherwise noted, ***p < 0.001, **p < 0.01, *p < 0.05.

**Figure 1:**
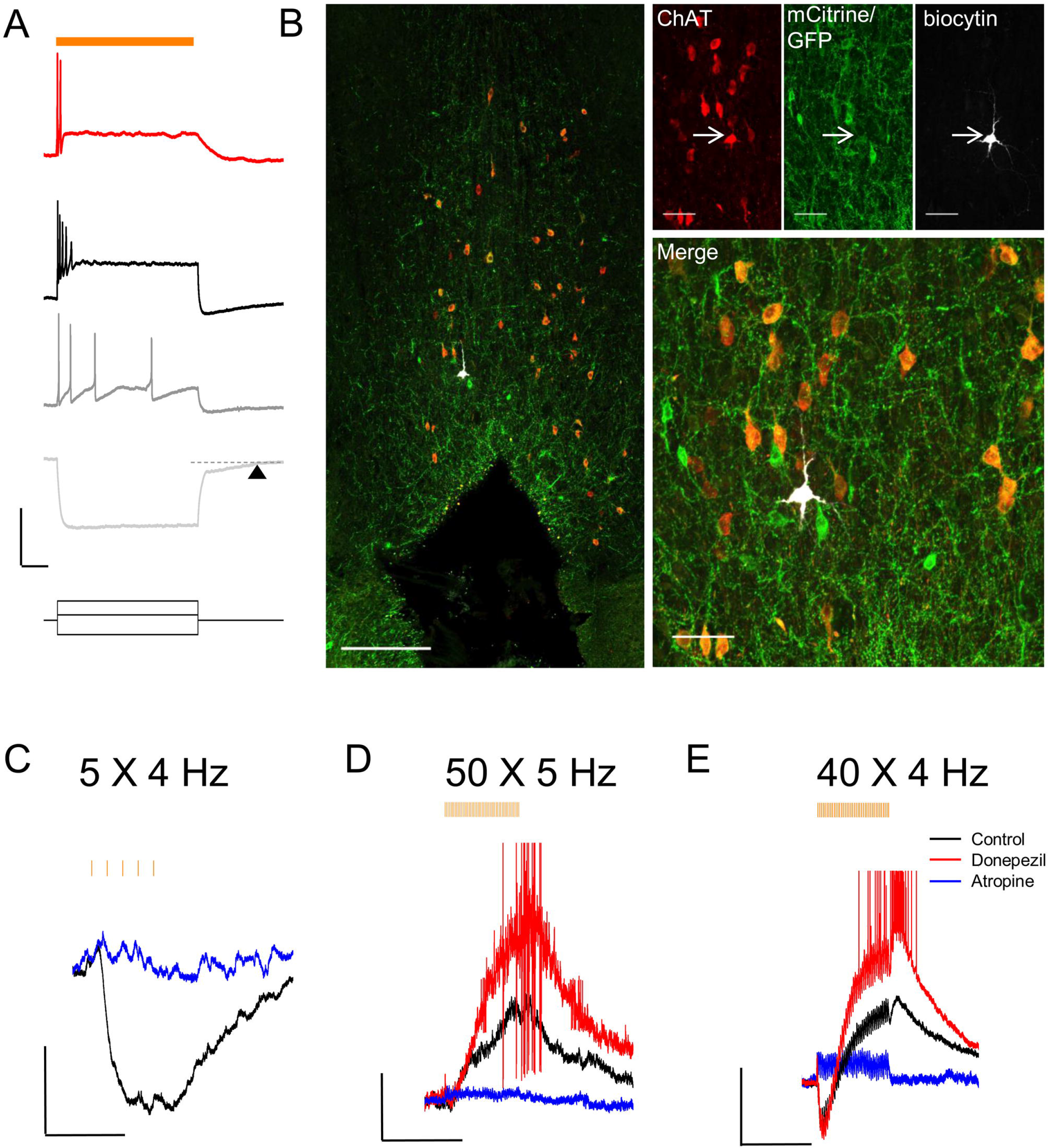
Red-shifted optogenetic protein (ReachR) was functionally expressed in MS/DBB cholinergic neurons and terminals of Chat-cre x ReaChR-mCitrine mice. **(A)** *Upper:* Representative trace (Red) showing a MS/DBB cholinergic neuron that depolarized and fired action potentials (APs) in response to a 600 ms long pulse of yellow light (indicated by orange bar); The membrane potential changes in the same cell in response to injection of −200 pA (Light Gray), +75 pA (Dark Gray) and +275 pA. The time delay in returning to baseline (dashed line) following hyperpolarization in response to a negative current injection is indicated by the black triangle; Scale bars: x = 200 ms and y = 20 mV. **(B)** *Left:* Image of the MS/DBB in a coronal brain slice: anti-choline acetyl transferase (ChAT, red); anti-GFP antibodies (green); biocytin filled cell (white) from (A). *Right:* magnified image of the same cell (left) showing the biocytin filled cell was positive for ChAT (red) and GFP (green). **(C-E)** CA1 interneuron responses to 4 or 5 Hz optogenetic stimulation displaying **(C)** hyperpolarization, **(D)** depolarization or **(E)** biphasic responses; scale bars: x = 1000 ms and y = 5 mV.

**Figure 2:**
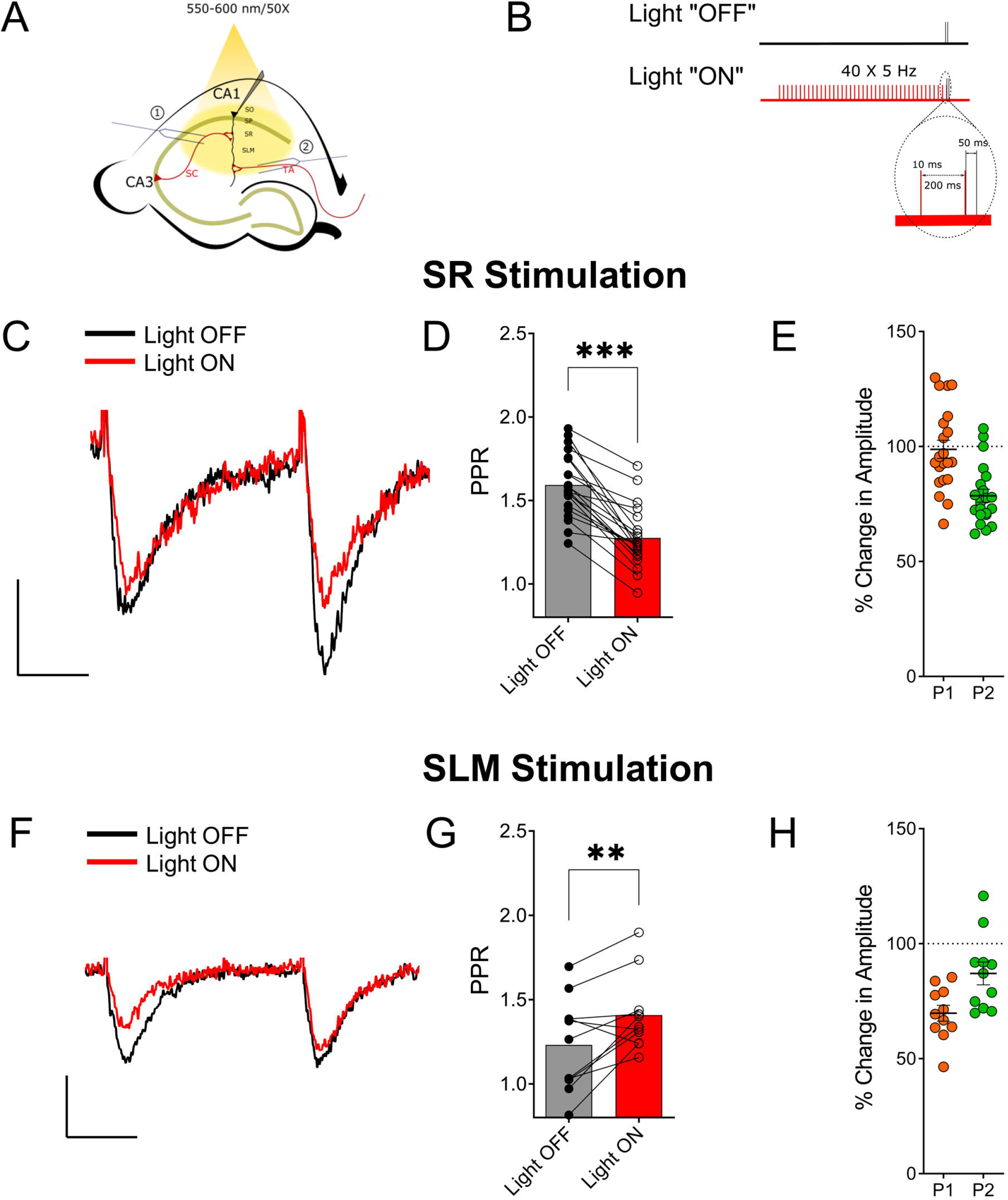
Optogenetically released acetylcholine (ACh) had different effects on Schaffer collateral (SC) and temporoammonic (TA) synaptic inputs. **(A)** Schematic representation of the experimental configuration: the bipolar stimulating electrode was placed either in (1) the stratum radiatum to stimulate the SC or (2) the stratum lacunosum moleculare to stimulate the TA pathways. **(B)** Schematic representation of the experimental paradigm; inset shows magnified view of the region marked by dashed line in the “Light ON” condition to show the temporal relationship between the last 2 light pulses (in red) and the paired electrical pulses (in black). **(C)** Representative paired-pulse traces from a CA1 pyramidal cell in response to SC stimulation with (red) and without (black) optogenetic stimulation of ACh release. **(D)** Scatter plot of individual data point overlaid on bar plot shows the effect of ACh release on SC PPR (paired t-test, p < 0.001). **(E)** Scatter plot displaying the %-change in amplitude in the first pulse (P1) and second pulse (P2) in response to ACh release before SC stimulation. **(F)** Representative paired-pulse traces from a CA1 pyramidal cell in response to TA stimulation with (red) and without (black) optogenetic stimulation of ACh release. **(G)** Scatter plot of individual data point overlaid on bar plot to show the effect of ACh release on TA PPR (paired t-test, p = 0.0096) **(H)** Scatter plot displaying the %-change in amplitude in the P1 and P2 in response to ACh release before TA stimulation; Scale bars: x = 25 ms; y = 50 pA.

**Figure 3:**
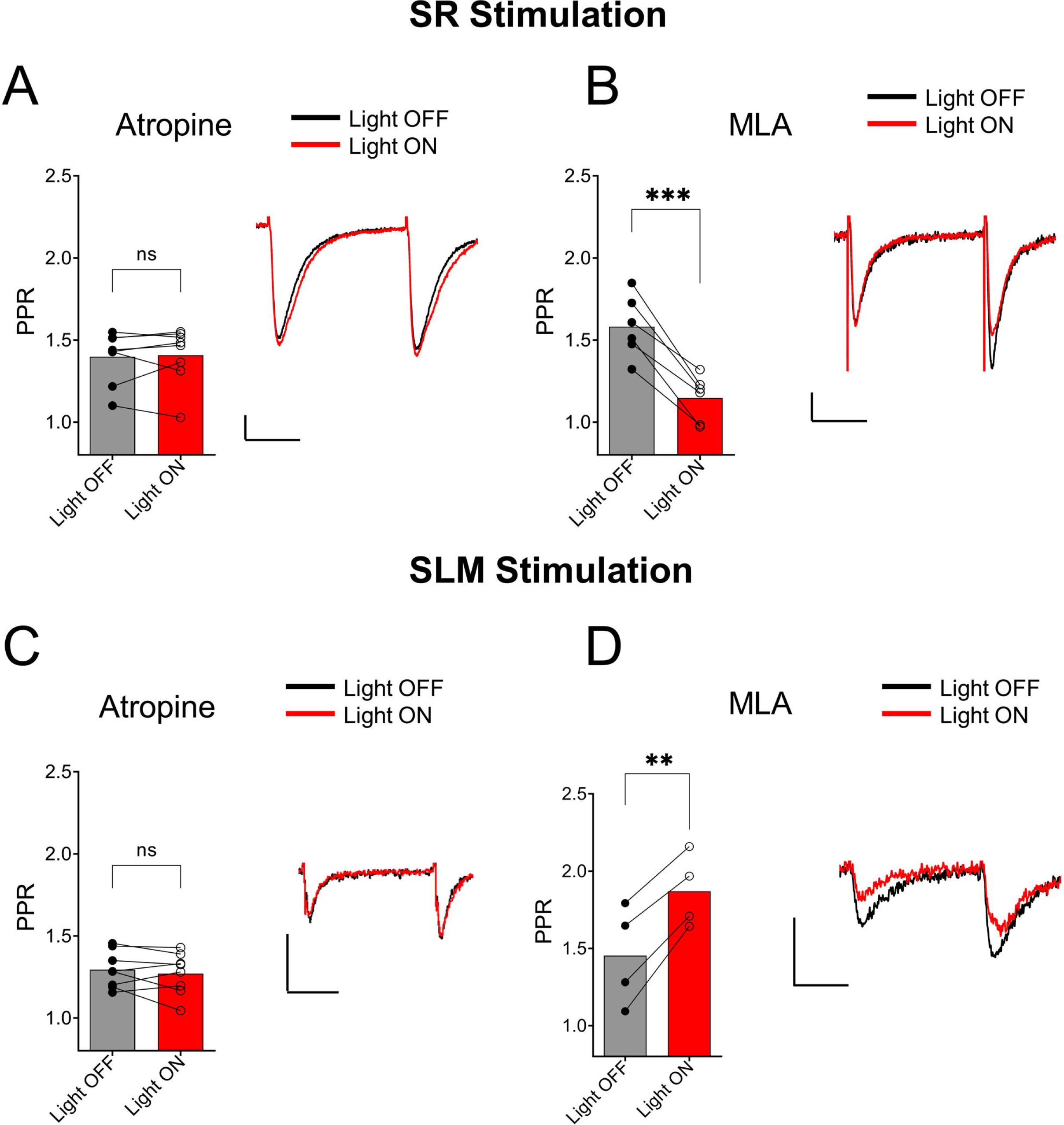
Muscarinic receptor activation mediated synaptic inhibition by ACh release in both the SC and TA pathways. **(A)** Bar plot of SC PPR (left) and representative EPSCs (right) demonstrates that atropine prevented ACh (red) mediated suppression of SC synaptic amplitudes and PPR (paired t-test; p = 0.7629). **(B)** Bar plot of SC PPR (left) and representative EPSCs show MLA had no effect on ACh (red) mediated inhibition of SC EPSCs and PPR (paired t-test; p = 0.0007). Scale bars: x = 20 ms; y = 50 pA **(C)** Bar plot of TA PPR (left) and representative EPSCs (right) show atropine blocked both the ACh (red) mediated inhibition of TA EPSC amplitudes and increase in TA PPR (paired t-test; p = 0.7159). **(D)** Bar plot of TA PPR (left) and representative EPSCs (right) show MLA did not prevent the effect of ACh release on TA EPSC amplitudes or TA PPR (paired t-test; p = 0.0036). Scale bars: x = 20 ms; y = 50 pA.

**Figure 4:**
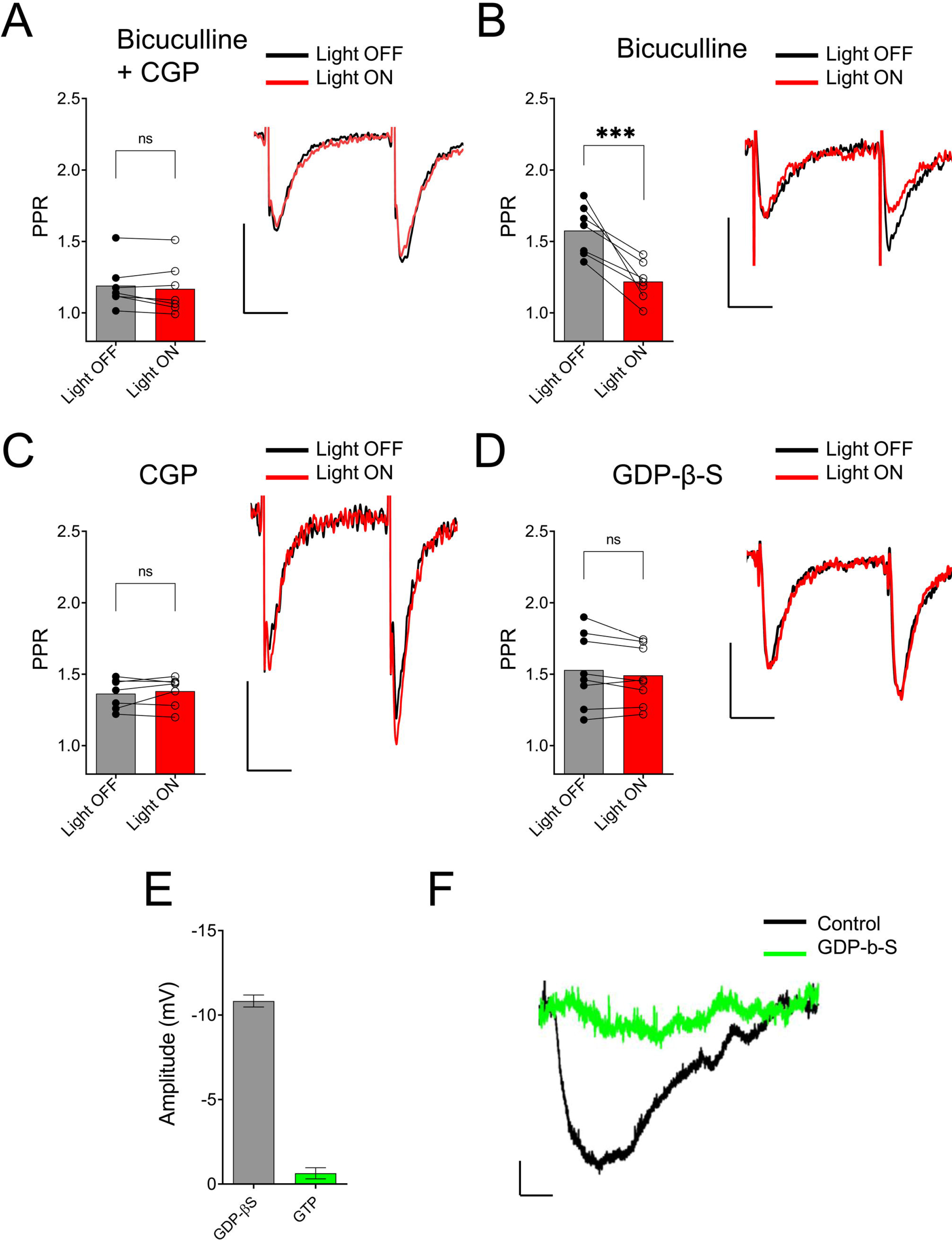
ACh-mediated inhibition of SC inputs required activation of postsynaptic GABAB receptors. **(A)** Bar plot (left) and representative EPSCs (right) showing ACh-mediated inhibition of SC EPSCs is prevented by GABAA antagonist bicuculline (BIC) and GABA_B_ antagonist CGP 52432 (Paired t-test; p = 0.2440); **(B)** Bar plot (left) and representative EPSCs (right) showing BIC alone had no effect on ACh-mediated inhibition of SC EPSCs (Paired t-test; p = 0.8332); **(C)** Bar plot (left) and representative EPSCs (right) showing ACh-mediated inhibition of SC EPSCs is blocked by bath application of CGP 52432 alone (Paired t-test; p = 0.0023) **(D)** Bar plot (left) and representative EPSCs (right) demonstrate that inclusion of GDP-β-S in the patch pipette prevented ACh-mediated reduction of SC EPSCs (Paired t-test; p = 0.1589). Scale bars: x = 20 ms; y = 50 pA. **(E)** Bar plot showing that hyperpolarization produced in CA1 PCs by the GABA_B_ receptor agonist baclofen was prevented when including GDP-1-S in the intracellular solution (white bar). **(F)** Representative membrane potential responses to baclofen in PCs recorded with intracellular solution containing GTP (black) and GDP-1-S (green); scale bars: x = 100 s; y = 2 mV.

## Results

### Immunofluorescent and Functional Verification of ReaChR Expression in the MS/DBB Cholinergic Pathway

To optogenetically release ACh from MS/DBB cholinergic terminals in mouse hippocampal brain slices, we crossed a driver mouse line that expressed Cre-recombinase under the control of the choline acetyltransferase gene (Chat-Cre) (Rossi et al. 2011) to a reporter mouse line that contained a Cre-dependent coding sequence for a red-shifted designer channelrhodopsin (ReaChR-mCitrine) knocked into the Rosa26 locus (ReaChR-mCitrine Cre-reporter mice) (Hooks et al. 2015; Lin et al. 2013). The selective expression of ReaChR in MS/DBB cholinergic neurons was confirmed by measuring light-evoked depolarizations in mCitrine fluorescent neurons of MS/DBB brain slices using whole cell patch clamp techniques (Fig 1A, upper panel). With resting membrane potential near −60 mV, 600 ms duration light-evoked responses were often accompanied by 1-2 action potentials that occurred near the beginning of the light pulse. We further assessed the electrophysiological responses to hyperpolarizing and depolarizing current steps (Figure 1A, lower panel). Membrane potential hyperpolarizing responses displayed a delayed return to baseline membrane potentials (200-300 ms) after termination of the current pulse. Furthermore, MS/DBB cholinergic cells exhibited strong spike frequency adaptation in response to positive current injection steps. These observations were consistent with previous descriptions of cholinergic neurons in the MS/DBB (Brazhnik and Fox 1997; M. Wu 2004; Min Wu et al. 2004).

By including 0.2% biocytin in the intracellular patch pipette solution, we were able to post-hoc identify the cell from which we recorded. To confirm whether the recorded cell was cholinergic, brain slices were fixed and processed for immunofluorescence using an antibody against choline acetyltransferase (ChAT). Furthermore, because the endogenous mCitrine fluorescent signal was weak, it was amplified using an anti-GFP antibody conjugated to AlexaFluor 488. Analysis of the tissue revealed that the ChAT-positive soma in MS/DBB slices co-localized with the AlexaFluor 488-labeled cells. The recorded neuron stained positively for both ChAT and mCitrine (Fig. 1B).

We next confirmed that light stimuli released ACh in hippocampal slices. Light stimuli (4 or 5 Hz train of yellow light pulses) produced the expected depolarizing, hyperpolarizing or biphasic response types similar to what had been previously reported by us utilizing 8 or 20Hz stimulation frequency in CA1 interneurons (Fig.1.C-E) (Bell, Bell, and McQuiston 2015b; Lawrence et al. 2006; McQuiston and Madison 1999c). Furthermore, the cholinesterase inhibitor donepezil (0.1 μM) increased response amplitudes, whereas the muscarinic receptor antagonist atropine (5μM) inhibited the responses. Some cells (Fig. 1E) responded with complex waveforms in which faster atropine-resistant responses (likely mediated by nicotinic ACh receptors) were superimposed on the atropine-sensitive slower waveform. Based on these observations, we concluded that the animal cross was suitable for studying the effects of ACh release in hippocampal slices.

### Optogenetic Release of ACh had Differing Effects on PPR on Input in the SR and SLM

Previous studies have reported that activation of presynaptic muscarinic receptors causes inhibition of the SC and TA inputs (Fernández de Sevilla and Buño 2003; M. Hasselmo and Schnell 1994; Thorn et al. 2017). Alternatively, activation of nicotinic receptors has been shown to enhance glutamate release and reduce PPR in the SC (Maggi et al. 2003). However, immediate effects of the release of ACh on presynaptic cholinergic receptors has only been investigated for muscarinic receptors on SC inputs (Buño, Cabezas, and de Sevilla 2006). Therefore, using physiologically-relevant optogenetic stimulation rates of MS/DBB cholinergic axon terminals (>4 Hz, (Pascale Simon 2006; Hao Zhang, Lin, and Nicolelis 2011)) in hippocampal CA1, we investigated the effect of ACh release on glutamatergic neurotransmission in the SR and SLM. To do this we measured the amplitudes and PPRs of excitatory postsynaptic currents (EPSCs) in CA1 PCs following stimulation of SR and SLM inputs. We compared EPSC amplitudes and PPRs measured under control stimulation (light OFF) with those following the optogenetic stimulation of the cholinergic terminals (light ON). In the “Light ON” condition, a 5 Hz train of 40 pulses of yellow light (10 ms pulse width) was delivered immediately before stimulating SR or SLM inputs with a pair of electrical stimuli (40-120 μs in duration and 50 ms interval). Figures 2A and 2B schematically illustrate our experimental paradigm. Optogenetic release of ACh prior to paired-pulse electrical stimulation of SR inputs reduced the amplitude of the second EPSC (Light OFF: −138.8 ± 11.95 Vs. Light ON: −108 ± 9.301; p < 0.0001), but had no significant effect on the first EPSC amplitude (Light OFF: −89.95 ± 9.104 Vs. Light ON: −86.47 ± 7.908; p = 0.3725) (Fig. 2C, E). This resulted in a significant reduction of the paired-pulse ratio in SR following ACh release (Fig. 2C-D; p < 0.001; t = 9.041; df = 20; n = 21 cells from 9 animals).

In contrast, optogenetic release of ACh prior to electrical stimulation of SLM inputs significantly reduced the amplitudes of both the first (Light OFF: −57.34 ± 9.002 Vs. Light ON: −38.93 ± 6.073; p = 0.0007) and the second EPSCs (Light OFF: −66.92 ± 9.239 Vs. Light ON: −56.53 ± 7.977; p = 0.0062) (Fig. 2 F, H). Furthermore, ACh release caused an increase in the PPR of inputs in the SLM (Fig. 2F-G; p = 0.0096; t = 3.194; df = 10; n = 11 cells from 6 animals).

Although most experiments were conducted in the presence of the acetylcholinesterase inhibitor donepezil (100 μM), a significant reduction in PPR in SR was also observed in donepezil’s absence (Light OFF: 1.662 ± 0.047 Vs. Light ON: 1.411 ± 0.045; p = 0.0029; t = 4.829; df = 6; n = 7 from 4 animals). However, we failed to see an effect of ACh release on PPR in SLM in the absence of donepezil (Light OFF: 1.278 ± 0.038 Vs. Light ON: 1.346 ± 0.058; p = 0.0618; t = 2.221; df = 7; n = 8 from 4 animals).Therefore, these results indicated that synaptically released ACh had distinct effects on glutamatergic neurotransmission in the SR and SLM.

### Muscarinic ACh Receptor Activation Mediates the Effect of ACh Release on Neurotransmission in the SR and SLM

We next investigated the type of cholinergic receptors that underlie the effect of ACh release on synaptic transmission in SR and SLM inputs. In the SR, the muscarinic receptor antagonist atropine (5 μM) blocked the ACh mediated reduction of PPR (Fig. 3A, p = 0.7629; t = 0.3137; df = 7; n = 8 cells from 3 animals). In contrast, inclusion of the α7 nicotinic receptor antagonist MLA (0.1 μM) in the extracellular solution had no effect (Fig 3B; p = 0.0007; t = 7.418; df = 5; n = 6 cells from 3 animals). Similarly, in the SLM atropine blocked the ACh-mediated increase in PPR (Fig. 3C, p = 0.7159; t = 0.3791; df = 7; n = 8 cells from 3 animals), whereas MLA had no effect (Fig. 3D; p = 0.0036; t = 8.368; df = 3; n = 4 cells from 3 animals). Therefore, release of ACh inhibited glutamatergic inputs in the SR and SLM via the activation of muscarinic cholinergic receptors.

### Activation of Postsynaptic GABAB Receptors Mediate the Effect of ACh Release on SC Neurotransmission

One possible mechanism to explain the ACh-mediated effect on PPR in the SR is through an indirect increase in the excitability of CA1 GABAergic interneurons (McQuiston and Madison 1999b; Parra, Gulyás, and Miles 1998; Pitler and Alger 1992). To test this possibility, we inhibited GABA_A_ and GABA_B_ receptors through bath application of 25 μM bicuculline (BIC) and 2 μM CGP 52432, respectively. As shown in Figure 4A, bath application with BIC and CGP 52432 blocked the ACh-mediated reduction of PPR (p = 0.2440; t = 1.292; df = 6; n = 7 cells from 4 animals). To determine the relative contribution of these two GABA receptor subtypes, we repeated the experiments with either BIC or CGP 52432 alone. Bath application of CGP 52432 alone was sufficient to block the ACh mediated reduction of PPR (Fig 4C; p = 0.8332; t = 0.2186; df = 7; n = 8 cells from 4 animals). In contrast, ACh release in the presence of BIC resulted in a reduction in PPR (Fig. 4B; p = 0.0023; t = 5.054; df = 6; n = 7 cells from 3 animals). Therefore, these data suggest that the optogenetic release of ACh increased the excitability of interneurons and suppressed the pathway in SR by the activation of GABA_B_ receptors.

GABA_B_ receptors are expressed on both presynaptic SC terminals and postsynaptic CA1 PCs. Thus, GABA release could inhibit synaptic transmission in SR through the activation of GABA_B_ receptors on pre- and/or postsynaptic membranes. We first investigated the possibility that ACh release increased the excitability of interneurons that activate postsynaptic GABA_B_ receptors on CA1 PCs. To do this, we performed experiments with an intracellular recording solution that contained 5 μM GDP-β-S, (instead of GTP) to inhibit postsynaptic G-protein receptor signaling (Bell, Bell, and McQuiston 2015b). As shown in Figure 4D, the inclusion of GDP-β-S in the patch pipette blocked the ACh mediated reduction of PPR (p = 0.1589; t = 1.576; df = 7; n = 8 cells from 5 animals). Intracellular GDP-β-S also inhibited baclofen (10 μM) mediated hyperpolarization of CA1 PCs demonstrating its ability to uncouple G-protein signaling (Fig. 4E and F; p < 0.0001; unpaired t-test; t = 19.67 df = 8; n = 6 cells for GDP-β-S, and 4 cells for control). Therefore, these data suggested that ACh-mediated reduction of PPR in the SR was mediated through the activation of postsynaptic GABA_B_ receptors.

### GABAB-mediated inhibition of excitatory synaptic transmission in SR is mediated by the activation of Inwardly Rectifying Potassium Channels

Because postsynaptic GABA_B_ receptors can mediate some of their actions through the activation of G-protein coupled inwardly rectifying potassium channels (GIRKs) (Lüscher et al. 1997), we tested whether GIRK activation mediated the ACh reduction of PPR in the SR. As a first test, we held CA1 PCs at hyperpolarized holding potentials to remove the intracellular blockade of GIRK channels by Mg^2+^ or polyamines. When we clamped the neurons at −90 mV, a significantly greater reduction of the PPR was observed (Fig. 5A; p = 0.0011; t = 4.731 df = 9; n = 6 cells held at −90 mV, and 5 cells held at −70 mV). Next, we attempted to inhibit the GIRK conductance by replacing K^+^ with Cs^+^ in the intracellular recording solution. The intracellular solution also contained 10 mM QX-314, a blocker of voltage activated Na^+^ channels and GIRK channels (Andrade 1991). Under these conditions the effect of ACh on PPR in the SR was abolished (Fig. 5B, p = 0.0758; t = 1.899; df = 16; n = 17 cells from 6 animals). To further test the involvement of the GIRK channels, we attempted to directly inhibit the GIRK conductance via extracellular application of Ba^2+^ (200 μM) (Breton and Stuart 2017). As shown in Figure 5C, extracellular application of Ba^2+^ blocked the ACh-mediated reduction of PPR (Fig. 5C; P = 0.4984; t = 0.7296; df = 5; n = 6 cells from 3 animals). Therefore, our data suggest that ACh release increases the excitability of inhibitory interneurons that activate GIRK channels on CA1 PCs through the activation of GABAB receptors.

**Figure 5:**
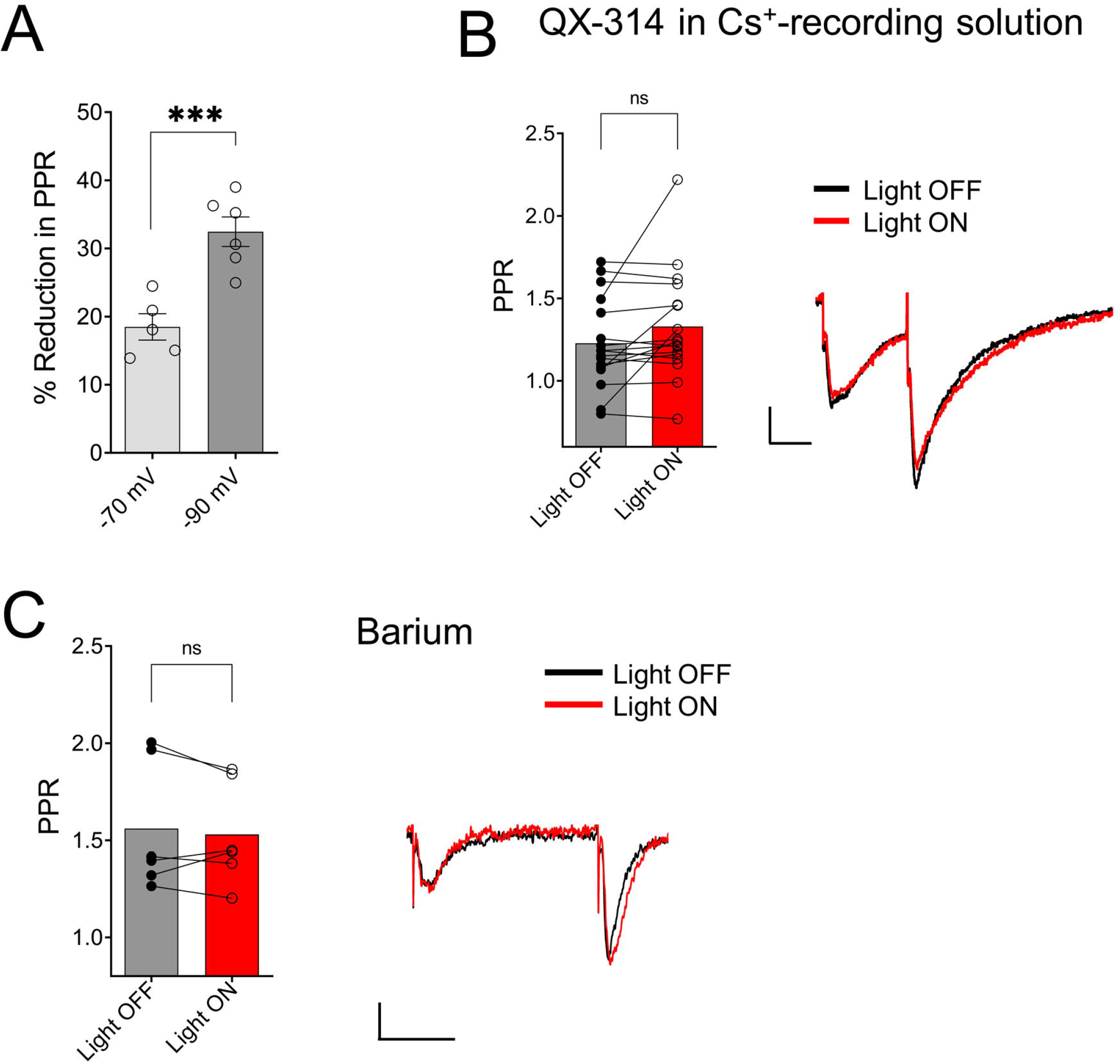
ACh-mediated reduction of SC EPSCs required activation of G-protein coupled inwardly rectifying potassium channels (GIRK) **(A)** Scatter plot of individual data points overlaid on bar plot demonstrating the ACh mediated reduction in SC EPSCs was larger at more negative holding potentials (unpaired t-test; p = 0.0011). **(B)** Bar plot (left) and representative EPSCs (right) shows inclusion of QX-314 and cesium in the intracellular recording solution prevented inhibition of SC EPSCs by ACh release (Paired t-test; p = 0.0758). **(C)** Bar plot (left) and representative EPSCs (right) shows extracellular barium prevented inhibition of SC EPSCs by ACh release (Paired t-test; p = 0.4984); Scale bars: x = 25 ms; y = 50 pA.

### ACh release inhibits synaptic transmission in excitatory inputs in the SR and SLM via the activation of different muscarinic receptor subtypes

Muscarinic receptors can be categorized into 2 groups, excitatory muscarinic receptors (M_1_, M_3_, and M_5_) that are coupled to G_q/11_ type G-proteins and inhibitory muscarinic receptors (M_2_ and M_4_) that have been shown to inhibit transmitter release via the activation of G_i/o_ type G-proteins (Levey and Edmunds 1995). Because the ACh-mediated inhibition of synaptic transmission in SR appears to involve an increase in inhibitory interneuron excitability, we tested the involvement of the M_1_ and M_3_ muscarinic receptors. To do this, we included the M_1_-selective antagonists VU 0255035 (10 μM) or the M_3_ receptor antagonist 4-DAMP (100 nM) in the bath. As shown in Figure 6A, the inclusion of VU 0255035 in the perfusate did not have an effect on the ACh-mediated reduction of PPR (p = 0.0061; t = 4.136; df = 6; n = 7 from 3 animals). In contrast, 4-DAMP blocked the effect of ACh on synaptic transmission in SR (Fig. 6B; p = 0.2209; t = 1.449; df = 4; n = 5 from 3 animals). Therefore, these data suggest that ACh release inhibited synaptic transmission in the SR through an increase in interneuron excitability via the activation of M_3_-muscarinic receptors.

**Figure 6:**
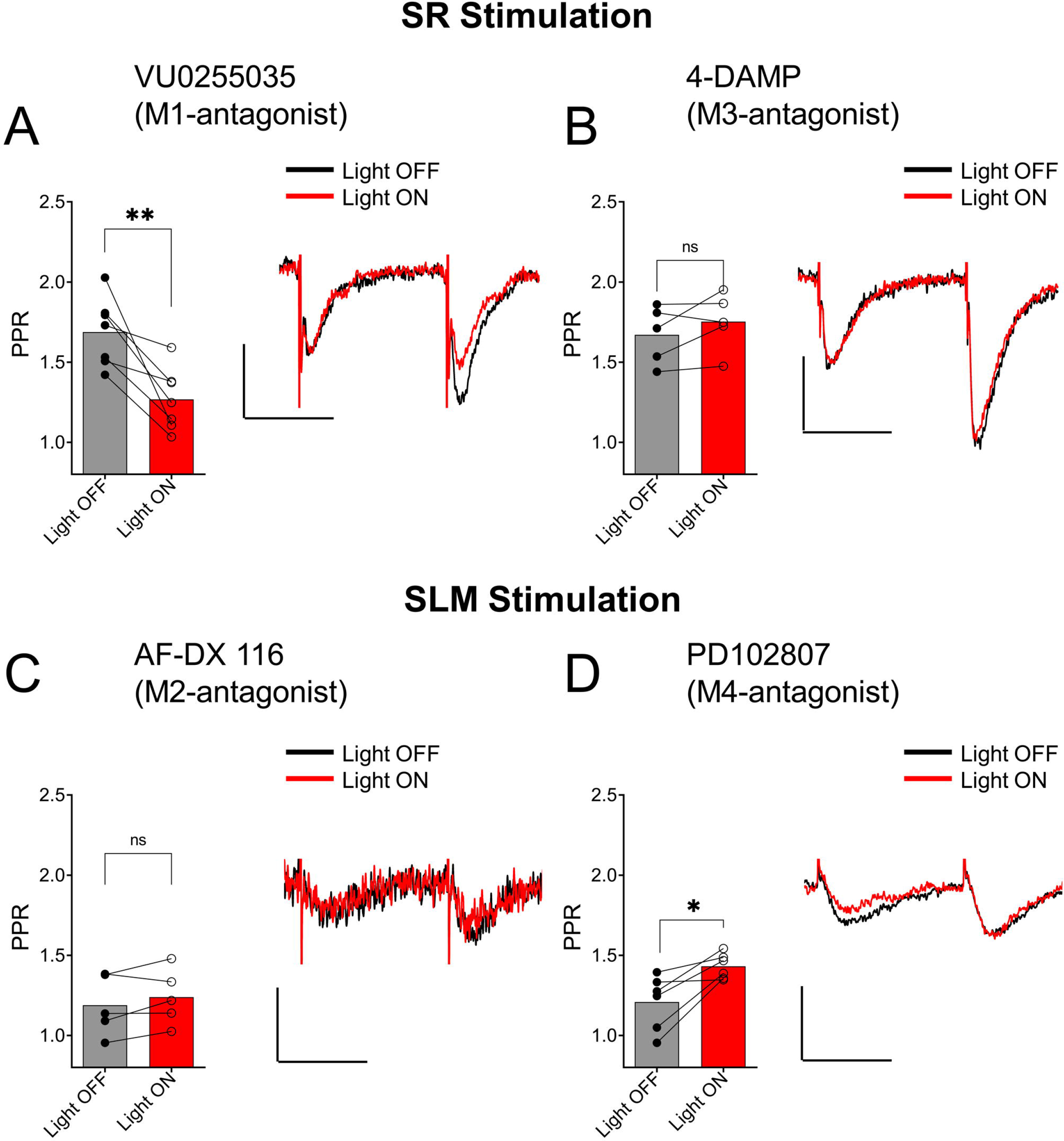
Distinct subtypes of muscarinic receptors mediated the inhibition of SC and TA EPSCs by ACh release. **(A)** Bar plot (left) and representative EPSCs (right) demonstrate that VU0255035 (M1 antagonist) had no effect on inhibition of SC EPSCs by ACh release (Paired t-test; p = 0.0061). **(B)** Bar plot (left) and representative EPSCs (right) demonstrate inhibition of SC EPSCs by ACH release is prevented by 4-DAMP (M3 antagonist) (Paired t-test; p = 0.2209). **(C)** Bar plot (left) and representative EPSCs (right) demonstrate that AF-DX 116 (M2 antagonist) prevented ACh-mediated inhibition of TA EPSCs (Paired t-test; p = 0.1984). **(D)** Bar plot (left) and representative EPSCs (right) demonstrate PD 102807 (M4-antagonist) had no effect on TA EPSC suppression by ACh release (Paired t-test; p = 0.0138); Scale bars: x = 20 ms; y = 50 pA.

We next investigated whether inhibition of excitatory inputs in the SLM was mediated by M_2_ or M_4_ muscarinic receptors. Application of the M_2_ receptor antagonist AF-DX 116 (500 nM) significantly blocked the ACh-mediated inhibition of synaptic transmission (Fig. 6C; p = 0.1984; t = 1.540; df = 4; n = 5 from 3 animals). In contrast, antagonism of the M_4_ receptor by PD102807 (1 μM) had no effect on the PPR (Fig. 6D, p = 0.0138; t = 3.711; df = 5; n = 6 from 3 animals). Therefore, these data suggest ACh release inhibited synaptic inputs in the SLM through the activation of M_2_ receptors.

## Discussion

In the present study, we demonstrate that low-frequency optogenetic stimulation of cholinergic terminals suppressed glutamatergic neurotransmission in the SR and SLM via different cellular and network mechanisms. ACh release indirectly inhibited SR inputs by increasing the excitability of CA1 interneurons that postsynaptically activated GABA_B_ receptors and GIRK channels on CA1 PCs. Furthermore, the ACh increase in CA1 interneuron excitability was mediated by the activation of M_3_ muscarinic receptors. In contrast, ACh release inhibited inputs in the SLM through the activation of M_2_ muscarinic receptors, likely located on the presynaptic terminals. Thus, our data have uncovered a previously unrecognized network mechanism by which ACh release controls SC inputs in hippocampal CA1 PCs through the modulation of interneuron excitability.

Previous studies from our lab (K. A. Bell et al. 2011; L. A. Bell, Bell, and McQuiston 2013, 2015b, 2015a) and others (Alger, Nagode, and Tang 2014; Zhenglin Gu and Yakel 2011) have studied the effects of ACh release on hippocampal function by utilizing virally mediated expression of blue light activated channelrhodopsin variants in MS/DBB cholinergic neurons. However, this approach may have limitations as blue light might not penetrate sufficiently far into tissue so that some cholinergic terminals are not be activated by light flashes. Furthermore, intracranial viral transfection may not sufficiently transfect all cholinergic neurons that project to the hippocampus due to imperfect targeting of the MS/DBB. In the present paper, we have taken a different approach in which the excitatory optogenetic protein ReaChR (Lin et al. 2013) was selectively expressed in cholinergic neurons through crossing a Cre-dependent ReaChR reporter mouse line (Hooks et al. 2015) to a cholinergic Cre-driver mouse line (Rossi et al. 2011). This has two potential advantages. First, ReaChR’s activation spectrum is red-shifted so that longer wavelength light can be used to penetrate further into tissue and excite more cholinergic terminals in a brain slice. Second, most if not all of the cholinergic neurons that project to the hippocampus express ReaChR. Using this approach, we demonstrated that ReaChR-expressing neurons in MS/DBB could be activated by yellow light pulses. More importantly, optogenetic stimulation of cholinergic terminals in hippocampal brain slices produced responses in hippocampal CA1 interneurons previously reported by others using electrical (Widmer et al. 2006) and blue light optogenetic stimulation (L. A. Bell, Bell, and McQuiston 2013). Therefore, by using genetically modified mouse driver and reporter lines, we can reliably release ACh from MS/DBB cholinergic terminals in hippocampal CA1 brain slices.

A small number of studies have investigated cholinergic receptor modulation of excitatory inputs in the SLM of hippocampal CA1 (M. Hasselmo and Schnell 1994; Thorn et al. 2017). Our studies extended these studies by being the first to investigate modulation of synaptically released ACh on excitatory afferents in SLM. Consistent with studies utilizing bath application of cholinergic agonists, we observed that ACh release suppressed excitatory inputs in SLM (M. Hasselmo and Schnell 1994; Thorn et al. 2017). Furthermore, ACh release appeared to act presynaptically as the inhibition of afferents in SLM was accompanied by an increase in the PPR, which is frequently used to determine presynaptic function. We further extended previous studies by demonstrating that suppression of SLM inputs by ACh release was mediated by muscarinic M_2_ receptor activation. These latter findings were consistent with observations from a recent study that suggest that presynaptic inhibition at synapses in the SLM was not mediated by M_4_ muscarinic receptors (Thorn et al. 2017). Therefore, our data confirms and extends observation made by previous studies and suggest that ACh release from MS/DBB cholinergic terminals results in presynaptic inhibition of inputs in the SLM via the activation of M_2_ receptors.

Several studies have investigated cholinergic modulation of SR inputs in hippocampal CA1 (M. Hasselmo and Schnell 1994; Kremin et al. 2006; Valentino and Dingledine 1981). Most studies utilized exogenous activation of muscarinic receptors that resulted in presynaptic inhibition of glutamate release from excitatory terminals in SR (Dasari and Gulledge 2011; M. Hasselmo and Schnell 1994; Kremin et al. 2006; Valentino and Dingledine 1981; Sheridan and Sutor 1990). Other studies that have utilized electrical stimulation to release ACh have confirmed that endogenous ACh caused a presynaptic inhibition of CA1 SR inputs through the activation of muscarinic receptors (Fernández de Sevilla and Buño 2003). In our study, we also assessed the effect of ACh release on SR inputs in hippocampal CA1 using optogenetic stimulation. However, we observed no effect of ACh release on the first electrically-evoked SC EPSC in a pair of stimuli. In contrast, the second EPSC of the paired stimulation was suppressed by ACh release. The suppression of the second EPSC was prevented by blockade of postsynaptic G-protein function, GABA_B_ receptor antagonists, GIRK channel inhibition, or by M_3_ receptor inhibition. Thus, our data suggested that the primary effect of ACh release on SC inputs was to increase the excitability of a subset of interneurons via an M_3_ receptor-mediated mechanism. These interneurons through a combination of M_3_ receptor-mediated increase in excitability and feedforward excitation by SR inputs drove the interneurons to spike and release GABA onto GABA_B_ receptors of CA1 PCs. The GABA_B_ receptor activation then activated GIRK channels on CA1 PCs and inhibited SC inputs onto these same cells. Thus, these data have identified a novel mechanism by which ACh release affects SC synaptic inputs in the SR of hippocampal CA1.

Despite the role of GABA_B_ receptor activation of GIRK channels in suppressing SR inputs onto CA1 PCs, no outward current or conductance change accompanied the postsynaptic inhibition. This observation is difficult to reconcile with well described GABA_B_ inhibitory synaptic potentials (IPSPs) measured in CA1 PCs (Dutar and Nicoll 1988). However, this observation is consistent with the demonstration that GABA_B_ receptors form a tight complex with Rgs7, Gβ5 and GIRK2 channels in CA1 PC dendritic spines (Fajardo-Serrano et al. 2013). In contrast, GABA_B_ receptors are segregated from this complex in dendritic shafts, which suggests that in CA1 PC dendrites GABA_B_ activation of GIRK channels maybe primarily confined to the dendritic spines. Considering that GIRK2 channels appear to be necessary for GABA_B_ IPSP activation in CA1 PCs(Marron Fernandez de Velasco et al. 2017), it is possible that the GABA_B_-mediated postsynaptic inhibition of SR inputs measured in our studies was confined to the synaptic spine and conductance changes could not be measured at the soma. Thus, suppression of SR inputs in individual spines of CA1 PCs may provide a mechanism for selective inhibition of individual synapses on CA1 PCs.

However, our results differ from previous studies that have investigated muscarinic modulation of SC inputs in hippocampal CA1. These differences could arise for a number of possible reasons. First, because there is a non-uniform density of cholinergic afferents and acetylcholinesterase in hippocampal CA1 (Aznavour et al., 2002; Franklin and Paxinos, 2007), uniform exogenous application of cholinergic agonists may activate presynaptic muscarinic receptors that are not normally activated by endogenous ACh. This is supported by the observation that the concentration of acetylcholine in different layers of the hippocampus varies during increased acetylcholine release, which accompanies theta rhythms (H. Zhang, Lin, and Nicolelis 2010). Second, studies that observed presynaptic muscarinic receptor-mediated inhibition of the pathways in the SR following electrically released ACh were performed at 20-22 ⁰C, in the presence of the acetylcholinesterase inhibitor physostigmine, and a GABA_A_ receptor antagonist (Fernández de Sevilla and Buño 2003). In contrast, we performed our studies at 32 – 34 ⁰C either in the absence of any inhibitors or in the presence of the acetylcholinesterase inhibitor donepezil. Importantly, acetylcholinesterase isoforms are temperature sensitive and the acetylcholinesterase inhibitors physostigmine and donepezil have different affinities for the varying isoforms of acetylcholinesterase expressed in the hippocampus (Zhao and Tang 2002). Thus, the studies performed at colder temperatures in the presence of physostigmine may have permitted ACh release to diffuse farther from cholinergic synapses and activate presynaptic muscarinic receptors on terminals in the SR that under our conditions could not be activated. Thus, it is unclear whether the observations made in the presence of physostigmine and colder temperatures uncovered an effect of ACh release that normally occurs physiologically or only happens under conditions of acetylcholinesterase inhibition such as in the treatment of AD patients. Nevertheless, our results in the absence of acetylcholinesterase inhibitors and at more physiological temperatures suggest that ACh release affects excitatory inputs in the SR by increasing the excitability of inhibitory interneurons and facilitating feedforward inhibition, which activate GABA_B_ receptors and GIRK channels on CA1 PCs.

In addition to presynaptic inhibition by muscarinic receptors, previous studies have demonstrated the presence of α7 nicotinic receptors on terminals in the SR, which when activated by exogenous nicotinic agonists facilitate the release of glutamate (Ji, Lape, and Dani 2001; Maggi et al. 2003; Sola et al. 2006). Moreover, endogenous activation of presynaptic α7 nicotinic receptors has been implicated in the induction of synaptic plasticity at this synapse (Zhenglin Gu and Yakel 2011; Z. Gu, Lamb, and Yakel 2012). However, to our knowledge there have been no studies that demonstrate that ACh release can potentiate glutamate release from CA1 SC terminals on a timescale of an individual synaptic event. Our data suggest that ACh cannot potentiate the release of glutamate from SC or TA terminals in hippocampal CA1 when coupled with an individual presynaptic action potential. However, our data did not examine the possibility that presynaptic nicotinic receptors modify synaptic strength on a long term timescale such as that which occurs with long-term potentiation (Zhenglin Gu and Yakel 2011).

## Conclusion

In conclusion, our data have shown that optogenetically released ACh in mouse hippocampal slices differentially inhibited glutamatergic synaptic transmission onto CA1 PCs depending on the input. Inputs in the SLM of CA1 were presynaptically inhibited by ACh release onto M_2_ muscarinic receptors as previously demonstrated by others. In contrast, inputs in the SR of CA1 were not directly modulated by ACh release. Instead, ACh release increased the excitability of a subset of interneurons that synapse on CA1 PCs, through an M_3_ muscarinic receptor-mediated mechanism. This increased interneuron excitability facilitated feedforward inhibition that resulted in postsynaptic inhibition via GABA_B_ receptor activation of GIRK channels. Thus, ACh modulation of CA1 SC inputs would depend on the amount of ongoing activity of SC inputs and may occur at the level of an individual spine.

## Author Contribution Statement

PG designed experiments, acquired and analyzed data and wrote the manuscript. ARM performed experiments, conceived and directed the research, reviewed the data and wrote and edited the manuscript. All authors reviewed the manuscript. All authors read and approved the final manuscript.

## Acknowledgements

This work was supported by NIH grants R21AG055073 and R01MH107507 to ARM.

